# Occupancy-collection models: Towards bias-corrected modeling of species’ distributions using unstructured occurrence data from museums and herbaria

**DOI:** 10.1101/2021.01.06.425644

**Authors:** Kelley D. Erickson, Adam B. Smith

## Abstract

The digitization of museum collections as well as an explosion in citizen science initiatives has resulted in a wealth of data that can be useful for understanding the global distribution of biodiversity, provided that the well-documented biases inherent in unstructured opportunistic data are accounted for. While traditionally used to model imperfect detection using structured data from systematic surveys of wildlife, occupancy-detection models provide a framework for modelling the imperfect collection process that results in digital specimen data. In this study, we explore methods for adapting occupancy-detection models for use with biased opportunistic occurrence data from museum specimens and citizen science platforms using 7 species of Anacardiaceae in Florida as a case study. We explored two methods of incorporating information about collection effort to inform our uncertainty around species presence: (1) filtering the data to exclude collectors unlikely to collect the focal species and (2) incorporating collection covariates (collection type and history of previous detections) into a model of collection probability. We found that the best models incorporated both the background data filtration step as well as the incorporation of collector covariates associated with the probability of collection. We found that month, method of collection and whether a collector had previously collected the focal species were important predictors of collection probability. Efforts to standardize meta-data associated with data collection will improve efforts for modeling the spatial distribution of a variety of species.

## INTRODUCTION

Understanding the extent of species ranges is a fundamental concern with many applications, whether it is establishing areas of conservation concern (Brooks et al. 2004), identifying the spread of invasive species (Václavík and Meentemeyer 2012, Veran et al. 2016, Briscoe et al. 2019), or predicting how species may respond to global change (Zurell et al. 2016, Scherrer et al. 2017, Graham et al. 2017, Requena-Mullor et al. 2019). Coupled with this growing urgency for establishing species’ ranges has been an explosion in the amount of occurrence data available due to the rise in the digitization of museum specimens as well as the growth of citizen science initiatives (Powney and Isaac 2015, Meineke et al. 2018). Originally collected for a wide variety of intents and purposes, there are striking geographic and temporal biases in when and where specimens are collected (Hijmans et al. 2000, Sólymos 2007, Meyer et al. 2016). Thus, much of our understanding of the distribution of life on Earth comes from so-called “unstructured” museum, herbarium, and citizen science data (in contrast to “structured” systematic sampling such as survey or atlas data).

To date, the most common method for utilizing unstructured data to recreate species’ distributions is to apply species distribution models (Peterson et al. 2011) (Figure 1a). Owing to the numerous sources of biases and inaccuracies associated with occurrence data (Hijmans et al. 2000, Sólymos 2007, Meyer et al. 2016), there have been several approaches to correct for these biases ranging from use of target-group background sites (occurrence records of other species likely to have been collected along with the focal species as a proxy of effort, Ponder et al. 2001), systematic sub-sampling in geographic or environmental space, the use of bias files based on proxies of search effort, and aggregating background points (Phillips et al. 2009, Boria et al. 2014, Fourcade et al. 2014, Varela et al. 2014, Vollering et al. 2019). In most of these approaches, collection bias is assumed to be “canceled” out by contrasting biased occurrence data with “filtered” background sites that are manipulated to have the same (presumed) bias as the occurrences. However, the actual bias in unstructured data is rarely observed. As a result, employing these methods is based on an assumption about the underlying bias.

**Figure 1.**
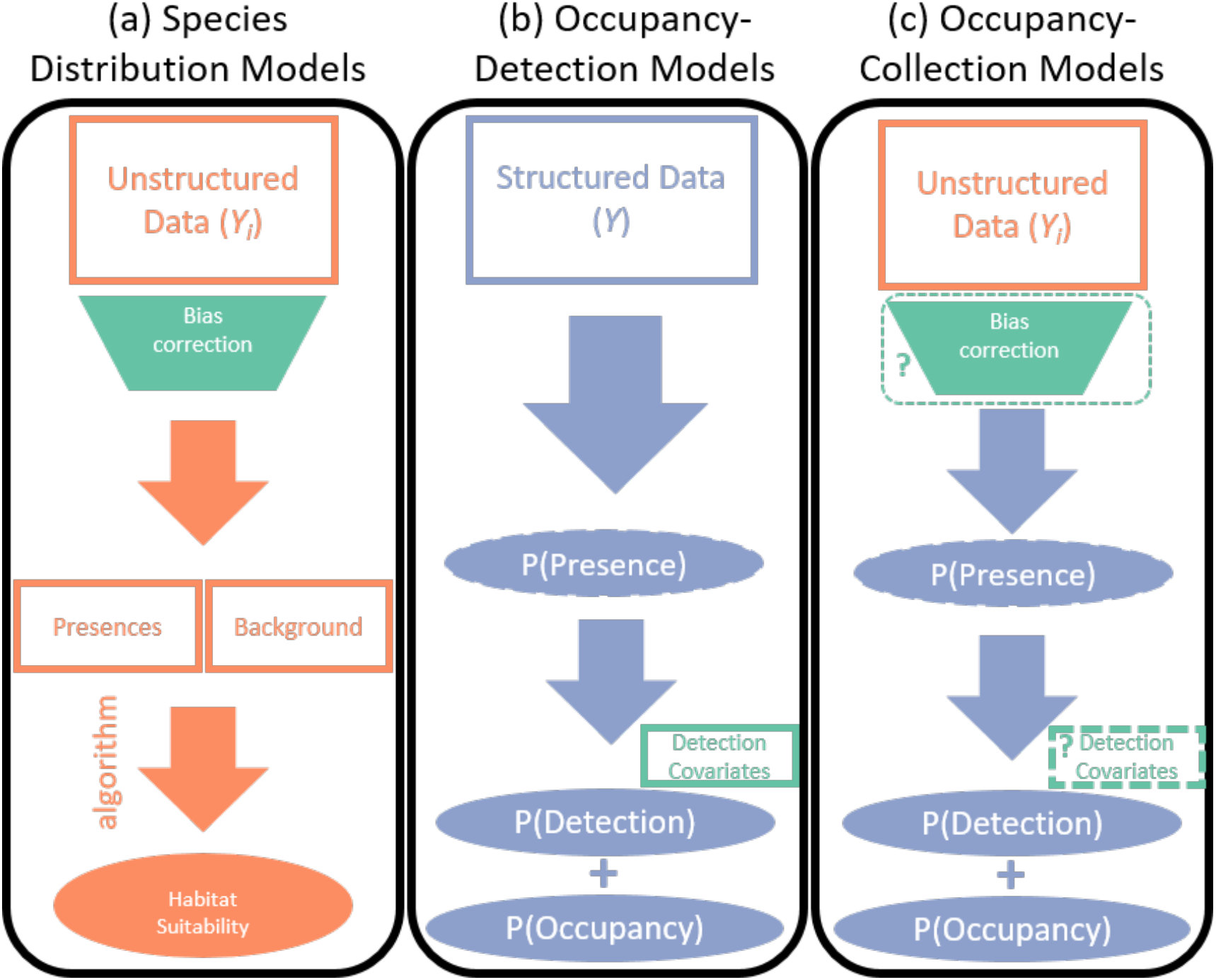
Schematic of modeling approaches for predicting species distributions: (a) species distribution models, which are frequently used to predict habitat suitability using unstructured occurrence data (i.e., museum or herbarium data), with little to no accounting for the processes underlying how the data was collected; (b) occupancy-detection models, which account for imperfect detection probabilities for systematic repeated-visit sampling; and (c) the modeling approach we introduce in this paper, which makes use of the unstructured occurrence data while accounting for the mechanism of collection. Colorblind-friendly colors from www.ColorBrewer.org by Cynthia A. Brewer, Geography, Pennsylvania State University.

**Figure 2.**
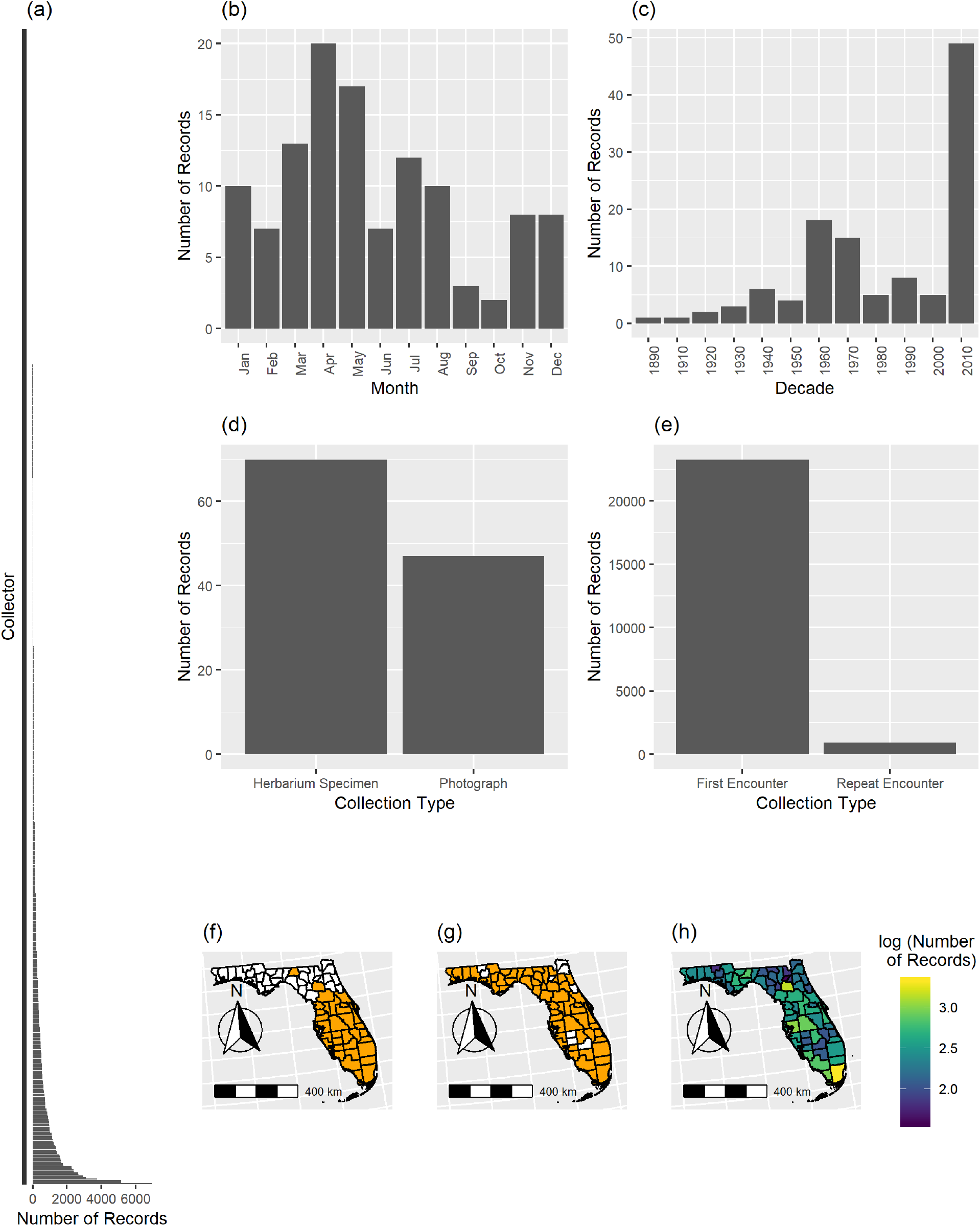
Biases in collection behavior. (a) Histogram of the number of records collected by each collector. Most collectors collect only one specimen. (b) Number of records of *Metopium toxiferum* collected in each month of the year. (c) Number of records of *M. toxiferum* collected in each decade. (d) Number of records of *M. toxiferum* that are deposited in an herbarium or a citizen science data portal. (e) Number of records of *M. toxiferum* that represent first encounters or repeat encounters. (f) Counties visited by the second-most prolific collector of specimens of Tracheophytes in Florida (O. K. Lakela). (g) Counties visited by the most prolific collector (J. R. Abbott). (h) Spatial map of collection effort (number of records of all species in the Tracheophytes) in each county in Florida on a log_10_ scale.

Originally developed for systematic repeated-visit wildlife surveys, occupancy-detection models (MacKenzie et al. 2002, 2003, 2018) explicitly account for the underlying processes which determine whether observers detect species (Figure 1b). The true occupancy status of a given site is an unknown latent state. When observers visit a site, the probability that they detect the focal species is a function of both the underlying reality of whether the site is occupied as well as the probability of detecting the focal species if it is present. Detection can depend on a number of covariates which can reflect, for example, time of year, observer experience, collection type (e.g., museum specimen, photograph), and observer identity. Occupancy-detection models have been increasingly used to analyze data from citizen-science projects, which often document aspects of search effort (e.g., number of observers, time spent searching) or ask that volunteers attempt to report all species of a known taxon (e.g., Bled et al. 2013, Van Strien et al. 2013, Berberich et al. 2016, Miller et al. 2016, Louvrier et al. 2018). These approaches assume that the data is structured in some way (i.e., involves defined surveying units with standardized instructions on how to collect and report data). Despite their utility for structured data, occupancy-detection models are challenging to apply to unstructured data because search effort is rarely documented (although see Wolf et al. 2011).

While the majority of digital occurrence data does not come from systematic sampling protocols, there is an associated wealth of often-overlooked data on how the specimen was collected on specimen labels that provides information about the collection process. Aside from simply identifying the species and where it was collected, specimen labels often also include the name of the collector and when it was collected. This type of information is commonly included in specimen databases, but its potential for informing bias correction when modeling species distributions has yet to be realized. Making use of these fields, it is possible to not only determine who collected a specimen but also recreate a collection history for a collector (e.g., Roberts et al. 2007) and thus collate unstructured data into a “structured” format for use in occupancy-detection modeling.

To use occupancy-detection models with unstructured occurrence data, careful decisions have to be made about how to translate it into a format that is appropriate for the occupancy-detection framework. One of the first considerations involves defining both the temporal and spatial scale of sampling. Typically, in studies involving structured survey data, whether they involve systematic wildlife surveys (e.g. MacKenzie et al. 2002) or atlas data (Bled et al. 2013, Altwegg and Nichols 2019), the sampling unit is determined prior to the initiation of the study. When using digital occurrence records, the definition of surveys must be made retrospectively. While it would be possible to use coordinate-points associated with occurrences to define sites, the vast majority of specimens are only able to be geolocated to much larger geo-political units, such as counties or equivalent administrative units(c.f., Collins et al. 2017, Park and Davis 2017, Pender et al. 2019). Estimating the probability of detection requires that sites are visited more than once (MacKenzie et al. 2002), so the spatial and temporal scale of sampling should be carefully chosen to ensure that each sampling unit has multiple occurrence records within it. At the other extreme, if the temporal scale is too large, the assumption that sites are closed to changes in occupancy within the surveying period may be violated, in which case, either the estimated probability of occupancy should be thought of as a “usage probability” instead of true occupancy (MacKenzie et al. 2018), or a dynamic “multi-season” occupancy-detection model should be used (MacKenzie et al. 2003). Checklists can be assembled by collating the occurrence records made by a unique collector within the retrospectively-defined survey window of space and time, and then pre-existing methods of translating checklist data into detection histories can be used (Altwegg and Nichols 2019). When constructing checklists, duplicate specimens of the same species within the sampling unit by the same collector should be discarded.

In this study, we address the question of how to account for biases in occurrence data when modeling species’ distributions. As a case study illustrating our methods, we focus on seven species in the family Anacardiaceae (cashews and sumac) in the US state of Florida. For each species, we collated metadata from occurrence records to reconstruct individual collectors’ collection histories, the type of occurrence record being made (herbarium sheet or photograph), and their taxonomic focus, as well as information on the timing of collection within and across years. Using this metadata, we were able to apply bias-correction frameworks from species distribution modeling and occupancy-detection modeling (Fig. 1), alone and in combination. More generally, we demonstrate the potential for using oft-overlooked information in collection records for accounting for bias in unstructured survey data. For simplicity, we focus here on the single-season occupancy-detection model (*sensu* MacKenzie et al. 2002), although our approach is amenable to any of the subsequent extensions of the basic occupancy-detection modeling framework that have been developed since then, such as dynamic multi-season models (MacKenzie et al. 2003), multi-species models (e.g. Van der Weyde et al. 2018), or models that address the issue of misidentification (Miller et al. 2011).

## MATERIAL AND METHODS

### Overview

We compiled occurrence record data for species in the family Anacardiaceae and for all Tracheophytes in the US state of Florida. For each species we constructed four models: (1) a full model with unfiltered background (non-detection) data, (2) a full model with filtered data, (3) a simple model with all data and (4) a simple model with filtered data. “Full” models incorporated covariates related to the probability of detection, whereas “simple” models did not. “Unfiltered” background data used data from all Tracheophytes as non-detections, whereas “filtered” data represented a subset of these records selected such that the collectors were more likely to have collected the focal species. We then evaluated the predictive accuracy of each model using cross-validation. We use the term “botanical collector” to refer to a person or team of people who collected a specimen in the field and deposited it into an herbarium, “citizen scientist” to a person who contributed a record in a citizen scientist data portal (e.g., iNaturalist), and “collector” to refer to either of these.

### Data downloads and processing

We downloaded occurrence records for all Tracheophytes in Florida from the Global Biodiversity Information Facility (GBIF download doi: https://doi.org/10.15468/dl.zsgv2l and https://doi.org/10.15468/dl.zsgv2l and https://doi.org/10.15468/dl.tzxjrm) on 12 and 13 February 2019. Although traditionally a repository for only museum and herbarium records, in recent years GBIF has included “research grade” records from citizen science data portals, which in our case, included iNaturalist (www.inaturalist.org), naturegucker (naturgucker.de) and Questagame (https://collections.ala.org.au/public/show/dr1902). County names and collector names were cleaned and filtered using regular expressions in R and openRefine (Huynh and Mazzocchi 2019). As only 42% of our downloaded occurrence records had coordinates, occurrence records with coordinates were converted to county-level records by determining which county polygon each point fell within based on the Database of Global Administrative Areas, version 2.8 (GADM; www.gadm.org). Botanical collectors often create duplicate specimen sheets from the same individual which are sent to other institutions. To remove these duplicates from our analysis, we kept only the first occurrence of a species by a particular collector on a particular date. Finally, we removed records that were missing information on who collected the record, which species was recorded, and when the record was recorded, leaving us with 166,230 county-level occurrence records of vascular plants in Florida. From these occurrence records, we selected all records for plants in the family Anacardiaceae, then of these retained only the seven species represented by ≥15 non-duplicate occurrences (Table A1 in Appendix A) for our focal species. We then modeled the distributions of these seven species.

Occurrence records were converted to detection histories as follows. First, records with identical collectors, months, years and county of collection were grouped together and denoted ‘collection event’ *t* (equivalent to a “survey” in the standard occupancy-detection framework; MacKenzie et al. 2002). For each collection event we defined

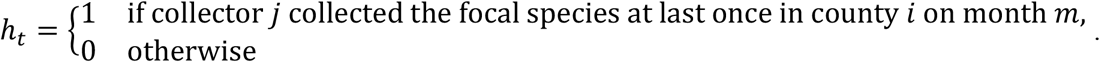

For each collector we examined their collection history across all their records and tallied the number of species and genera they collected, the months they collected in, and the number of each of our focal species they collected.

### Occupancy-Collection Models

We constructed a set of hierarchical Bayesian occupancy-detection models for each species. In each model, the ecological process of interest *Z*_*i*_, whether county *i* is occupied by the species of interest, is a Bernoulli random variable with probability *ψ*_*i*_:

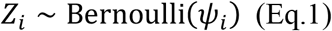

We assumed that the probability that a particular county was occupied depended on several covariates,

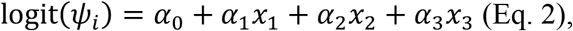

where *x*_1_ represented human population density; *x*_2_ average annual minimum monthly temperature; and *x*_3_ county area. Population density was obtained from Recht (2019) and minimum temperature from 2-arcmin PRISM climate data for 1895 through 2019 (Daly et al. 2002). All covariates were z-standardized (MacKenzie et al. 2018).

The occupancy status *Z*_*i*_ of site *i* is a latent state that we cannot directly measure. Instead, through the observation process we measure *y*_*i,t*_, whether a detection of the species occurs in collection event *t*. This observation process was modeled as:

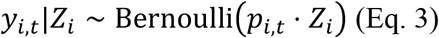

where *p*_*i,t*_ is the probability of detecting the species given that it is present. In the simple models, each county is assumed to have the same baseline probability of collection *p*, which has a flat beta prior. For the full models, the collection probability *p* depends on several collection-specific covariates,

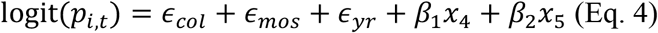

where *ϵ*_*col*_, *ϵ*_*mos*_, and *ϵ*_*yr*_ are random effects associated with collector, month and year, respectively. Since collectors are less likely to collect the same species in the same place twice (ter Steege et al. 2011), we defined a binary variable, *x*_4_, to designate whether or not a collector had previously collected the focal species in that county before. Variable *x*_5_ indicated whether the collector was a citizen scientist (*x*_5_ = 1) or an herbarium collector (*x*_5_ = 0). The full likelihoods and priors for both the simple and full model are available in Appendix B.

To evaluate the effectiveness of data filtration (Fig. 1a), we ran both the simple and full models first using the full set of detection histories (all data) and then using a subset of the data excluding collection events from collectors who only collected a single genus and never collected the focal species (filtered data).

For each of the four combinations (full model with all of the data, full model with filtered data, simple model with all of the data and simple model with filtered data), we ran two chains for 349000 iterations using the Nimble Bayesian modeling system version 0.9.0 (de Valpine et al. 2017) for R version 3.6.1 (R Core Team 2019), discarding the first 10000 iterations as burn-in. We assessed convergence using the Gelman-Rubin diagnostic (R-hat) and visually inspected the chains using the R package MCMCvis (Youngflesh 2018).

### Cross-validation

To assess the predictive accuracy of our models we used k-fold cross-validation with 5 folds for each of our species-model-data combinations. We assembled each fold by randomly dividing the list of counties into one of five groups. Since estimating random effects (Eq. 4) requires that there is at least one observation for each collector, year and month, if any of the resulting training data sets (assembled from the remaining folds) was deficient in any of these categories, one of the occurrence records of that level in the held-out fold was randomly selected and moved from the held-out data into the training data set. On each of the 5 runs, one fold was held out, the remaining training data was used to run the models, and the resulting model was used to predict the held-out data points. We evaluated the ability of each of the models to predict the held-out data using a scaled log-likelihood metric

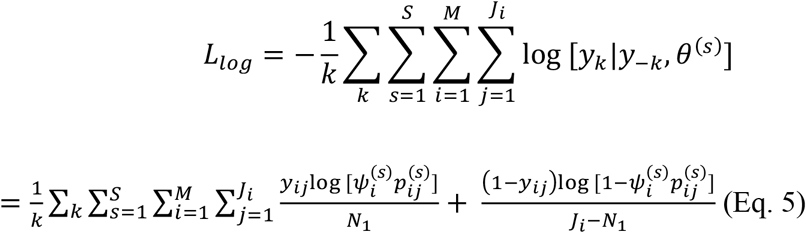

where *k* is the number of folds, *y*_*ij*_ is the held-out data for the *i* = 1, ⋯, *M* counties, and *t* = 1, ⋯, *J*_*i*_ surveys, where *J*_*i*_ represents the number of surveys of county *i* in the held-out data, evaluated over *S* total MCMC iterations. To allow comparison between the filtered and unfiltered models, which have different numbers of non-detections of the focal species, we scaled the respective likelihoods by the number of detections *N*_1_ and the number of non-detections (*J*_*i*_ − *N*_1_).

## RESULTS

### Collector Behavior

There were 762 people who collected at least a single Anacardiaceae species in Florida between the years 1895 and 2019, of whom 287 were citizen scientists and 475 herbarium collectors (Figure 3d). The number of records collected by a single collector was highly skewed, with some people collecting over 5000 records, while the most frequent number of records collected was one. There was temporal variation in the number of records collected by month as well as by decade (for example, as can be seen in Figure 3b and c for *M. toxiferum*). Most collection events represented the first time a collector collected that species in the county (Figure 3e). The spatial distribution of collections across Florida as a whole (Fig. 3h) and by collector, as can be seen, for example, with the two collectors who collected the greatest number of Tracheophytes in Florida (Figures 3f and 3g).

**Figure 3.**
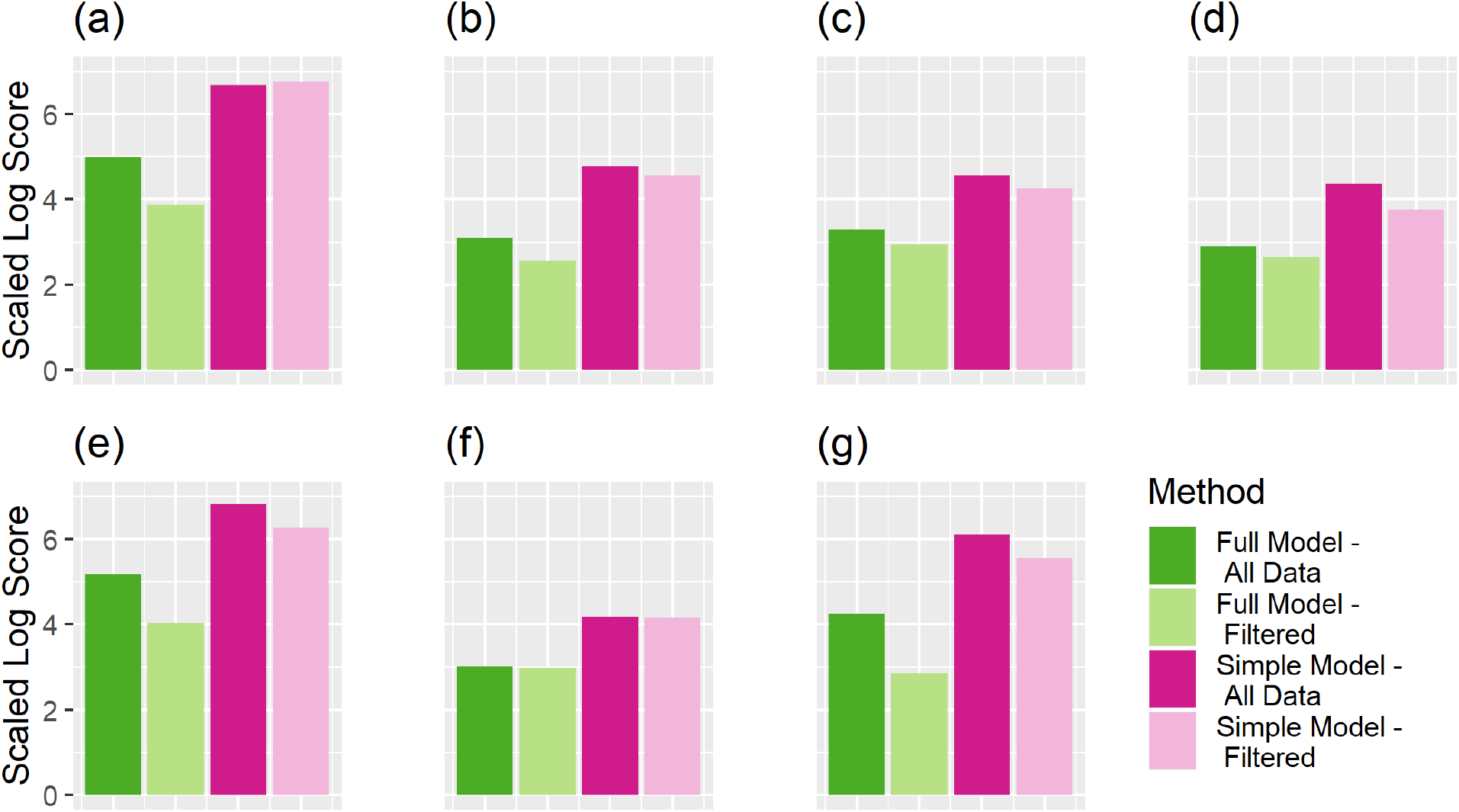
Cross-validation scores for the full models with all data (dark green), full models with filtered data (light green), simple models with all data (dark pink), and simple models with filtered data (light pink) for (a) *M. indica*, (b) *M. toxiferum*, (c) *R. copallina*, (d) *S. terebinthifolia*, (e) *T. pubescens*, (f) *T. radicans*, and (g) *T. vernix*. Smaller values indicate better predictive accuracy. Colorblind-friendly colors from www.ColorBrewer.org by Cynthia A. Brewer, Geography, Pennsylvania State University.

### Cross-validation

For each species, cross-validation demonstrated that the full model with collection covariates using filtered non-detection data had the highest predictive accuracy (lowest log-likelihood score) against withheld data, followed by the full model using all non-detection data, and then lastly the two simple models. For six of the seven species, the simple model using all of the data performed the worst (had the highest log-likelihood score, Figure 4). Each of the full models performed better than the simple models, while the magnitude of the effect of data filtration varied across the species, improving model performance for some species, while having little effect for others, indicating that including detection covariates improved predictive accuracy more than filtering the data.

**Figure 4.**
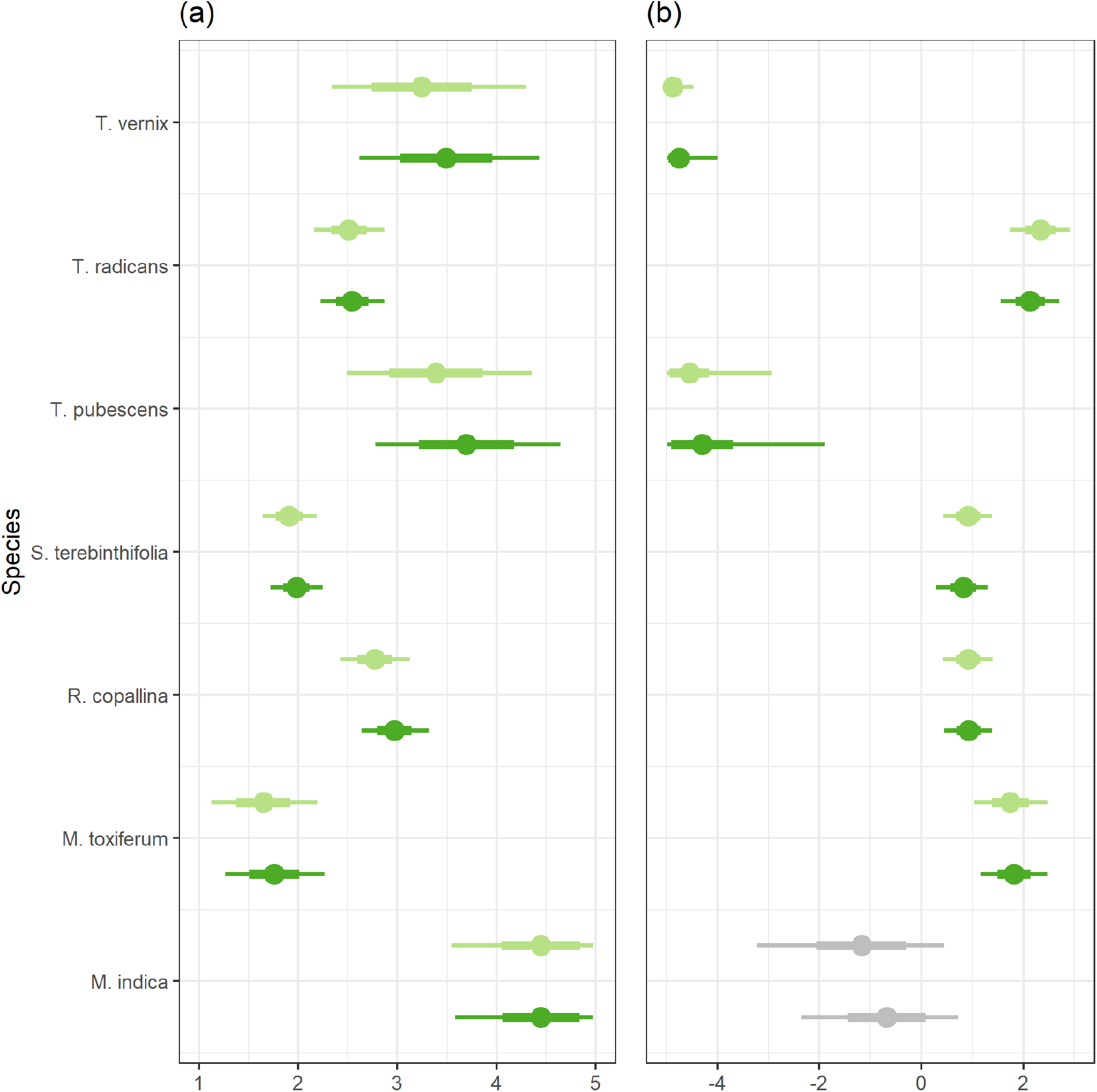
50% (thin line) and 95% (thick line) posterior credible intervals for (a) detection history and (b) type of collection (photograph or herbarium specimen). For each species, the bottom line (dark green) is the credible interval for the parameter estimate from the full model with all data and the top line (light green) is the credible interval for the parameter estimate from the full model with filtered data. Grey lines indicate that the 95% credible interval overlaps with zero. Colorblind-friendly colors from www.ColorBrewer.org by Cynthia A. Brewer, Geography, Pennsylvania State University.

### Detection covariates

For each species, both of the full models that included collector covariates indicated that if a collector had previously recorded the focal species, they were more likely to collect it again (odds ratios ranging from 5.21 to 85.29 more likely, odds ratios calculated as *e*^F^ where *μ* is the mean of the posterior distribution of the parameter). Citizen scientists had a higher probability of collecting *M. toxiferum, R. copallina, S. terebinthifolia*, and *T. radicans* (odds ratios from 2.50 to 10.38). Collectors who were preserving specimens on herbarium sheets were 3.23 and 94.28 times more likely to collect *M. indica* and *T. pubescens*, respectively, while *T. vernix* was only collected by those making herbarium specimens.

### Occupancy covariates

For each of the species, the four models mostly agreed with each other in terms of which covariates were important for predicting occupancy. The posterior density for the coefficient associated with population density overlapped zero for all of the models except the simple model using filtered data for *M. indica*, which suggested a positive association between human population density and occupancy. Minimum temperature was positively associated with occupancy for *M. indica, S. terebinthifolia*, and negatively associated with occupancy for *T. pubescens* and *T. vernix* for all four models for each species. County area was only relevant for one species, with all four models suggesting a positive relationship between county area and occupancy for *T. vernix*. (Appendix D Figure 1).

## DISCUSSION

The objectives of our study were to a) develop a framework for applying methods traditionally applied to structured data (occupancy-detection modeling); and b) evaluate whether bias-correction methods used in this framework and/or borrowed from species distribution modeling improved predictive accuracy. We found that models with the highest predictive accuracy incorporated both the background data filtration step inherited from the species distribution modeling framework (Fig. 1a) as well as the incorporation of collector covariates associated with the probability of collection from the occupancy-detection modeling framework (Figure 1b). Including collector covariates improved model performance more than data filtration (Figure 4).

We found that for all species, individuals who had previously collected the focal species had a much higher probability of collecting the species again (Figure 5a). Previous collection of the focal species is one way of measuring the experience of individual collectors, and several studies have indicated that observer experience is positively associated with increased detection probability (i.e., Berberich et al. 2016, Johnston et al. 2018), although previous work also suggests that individuals are more likely to collect species they have not previously collected (ter Steege et al. 2011). In our study, most collectors never collected the focal species (ranging from 58 to 96% of the 762 collectors).

**Figure 5.**
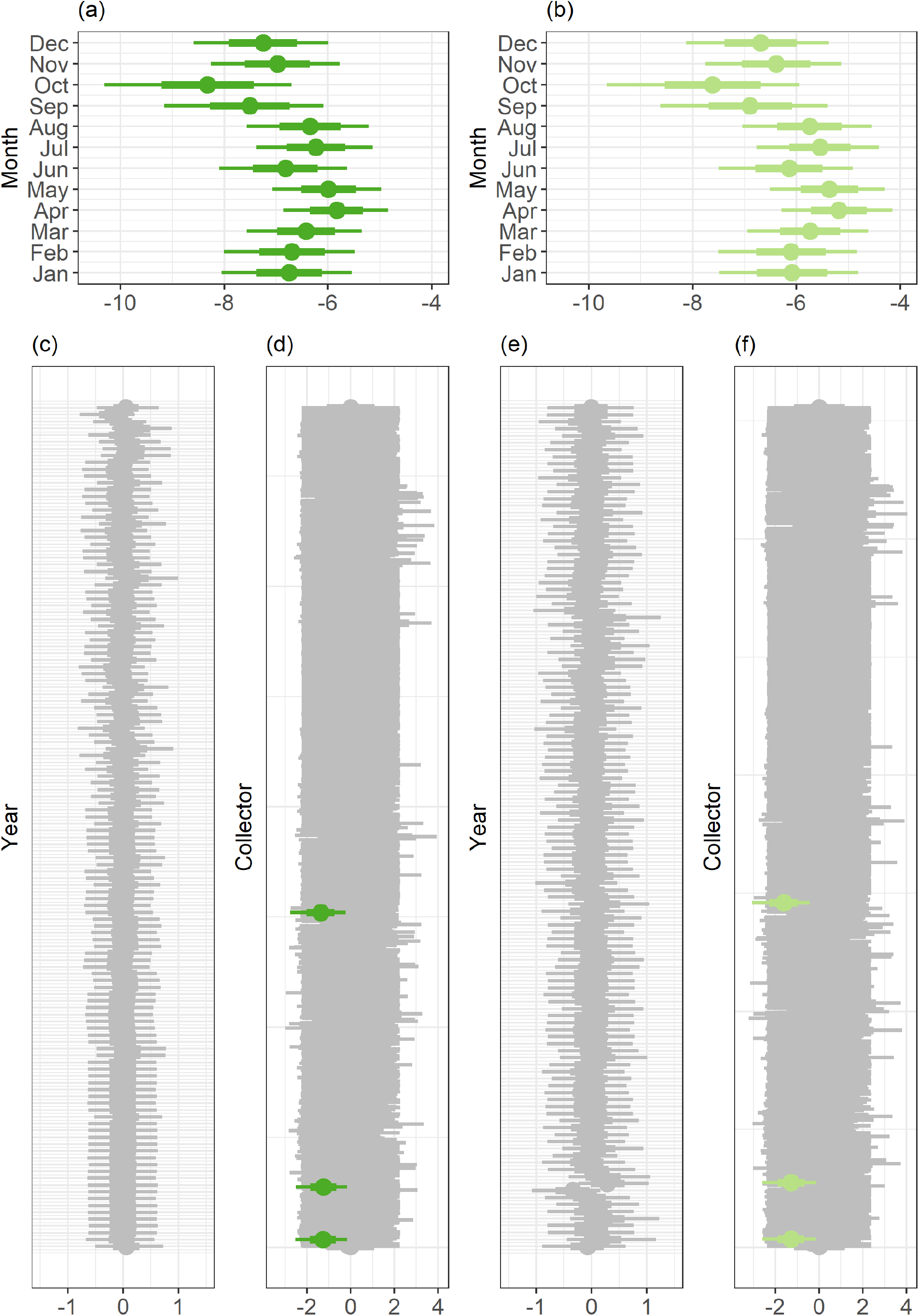
*M. toxiferum* 50% (thick line) and 95% (thin line) credible intervals for (a) month effect for the full model with all data, (b) month effect for the full model with filtered data, (c) collector effect for the full model with all data (d) year effect for the full model with all data (e) collector effect for the full model with filtered data, and (f) year effect for the full model with filtered data. Greyed-out credible intervals overlap 0. Colorblind-friendly colors from www.ColorBrewer.org by Cynthia A. Brewer, Geography, Pennsylvania State University.

We found species-specific differences in which type of collector (citizen scientists versus botanical collectors) was more likely to collect the focal species. Four of the species were more likely to be detected by citizen scientists (*M. toxiferum, R. copallina, S. terebinthifolia*, and *T. radicans*), while the remaining species were either never collected by citizen scientists (*T. vernix*) or more likely to be preserved as herbarium specimens *(M. indica* and *T. pubescens*). In contrast to the process by which a citizen scientist typically records a plant (which usually involves taking a photograph and possibly annotating it), preparing an herbarium specimen is a more involved process and involves physically obtaining the specimen, mounting it to a specimen sheet, and (for our uses), deposition of the label data into an electronic data portal. There are therefore possibly species-specific traits which could influence the probability of collection according to whether the collection was performed by a citizen scientist or botanical collector. Indeed, all of the members of the Anacardiaceae produce the secondary compound urushiol, a skin irritant (Judd et al. 2008), which could dissuade collectors who need to physically handle the specimens. Species also differ with regard to how easy they are to identify or even notice by non-experts (i.e., Tulloch et al. 2013, Bird et al. 2014, Meyer et al. 2016, Johnston et al. 2018).

There are well-documented temporal biases in digital occurrence data (Sólymos 2007, Meyer et al. 2016), that are also present in our data set (Figure 3c). However, while we found that collection probability did vary across the months for many of the species (Figure 3b, Appendix E), we did not see evidence of a year effect on collection, even for introduced species (Appendix E), which suggests that some of our other covariates are correlated with the year. For example, citizen science campaigns focused on biodiversity are generally a more recent phenomenon, so the rate of collection by citizen scientists is positively associated with year. Likewise, we also found little to no evidence that including a separate random effect for each collector was necessary (Figure 5, Appendix E), which also suggests that our existing covariates are adequately capturing the existing variation among collectors.

### Occupancy-collection modeling

Our work shows a clear advantage to modeling unstructured data with “structured” techniques which can explicitly account for bias. However, adapting occupancy-detection models for use with unstructured data requires careful consideration of model assumptions and requirements. These include populations closure, no un-modeled heterogeneity, how to handle false identifications, and choosing appropriate sampling units (e.g. MacKenzie et al. 2018, Altwegg and Nichols 2019). Depending on the time-scale of the study and the life-history of the organism, there is a tradeoff between strictly satisfying the closure assumption versus limiting the interpretation of the resulting occupancy estimate as a measure of habitat “usage” probability (MacKenzie et al. 2018). Carefully considering sources of variation in both detection and occupancy probabilities, as well as including random effects can help ensure that all sources of heterogeneity in the data is accounted for. The issue of false identifications has been well-documented in the citizen science literature (Ruiz-Gutierrez et al. 2016, Cruickshank et al. 2019), which can be addressed in the modeling process (Miller et al. 2011). Finally, a key decision of adapting unstructured data for use in an occupancy-detection framework requires careful consideration of how to define spatial units and surveys. We chose counties as our spatial units. However, it is possible that the same analysis could be replicated at smaller spatial scales (i.e., assigning records into grid cells). When determining the appropriate spatial scale to use and how to delineate what constitutes a survey, consideration should be made to ensure that the choice of these definitions results in each site having at least two independent visits.

We focused on collection covariates that were readily available and that we hypothesized could be relevant for all of the species we considered. In a smaller-scale study with fewer collectors, it would be possible to gather additional information about collectors that could provide a more complete picture. For example, the home institution of a given collector could provide additional information on where that person is most likely to collect. Leveraging information from additional sources such as the Index Herbariorum (http://sweetgum.nybg.org/science/ih/) and Bionomia (https://bionomia.net) that collate additional biographical information about collectors could allow for the incorporation of additional information to further distinguish among collectors. Harmonizing collector names across records and constructing collection histories is a laborious process. Mobilizing this store of information on collection would be aided by standardization of allowable entries in database fields, as well as advances in artificial intelligence and textual analysis.

### Conclusions

Our work is a test of how to account for widely-appreciated biases in unstructured data (Meyer et al. 2016) when modeling species’ distributions. There are numerous approaches to both modeling the distribution of species, and accounting for biases associated with the manner in which the data was collected. Species distribution models, which make use of the ever-increasing stream of digital biodiversity occurrence data, typically account for these biases through various data filtration steps but fall short of modeling the processes that lead to the resulting biases in the data (Figure 1a). On the other hand, occupancy-detection models explicitly model the role of imperfect detection in the data that we are able to collect, but they do so in a way that assumes the data come from structured repeat-visit surveys (Figure 1b). We found that utilizing both the bias-correcting filtration techniques from species-distribution modeling while explicitly modeling the manner in which the data was collected resulted in the best predictive accuracy. We therefore need methods like those explored in this study (Figure 1c), to allow for the use of digital occurrence data while explicitly accounting for the imperfect-detection processes that led to their existence.

## DATA AVAILABILITY STATEMENT

Upon acceptance, data will be deposited in an online repository.

## ACKNOWLEDGEMENTS

The authors thank Stephen Murphy, James Lucas, David Henderson, Matt Austin, and the Stephen Beissinger lab at UC Berkeley for helpful discussion and feedback.

