## Appendices for "Occupancy-collection models: Towards bias-corrected modeling of species’ distributions using unstructured occurrence data from museums and herbaria"

Appendix A.

Table 1: List of species of Anacardiaceae that have more than 30 records in GBIF and their introduced status (Wunderlin et al. 2019).

Table 1:

| Species | Common   Name | Status | Number   of Collectors | Number   of Records | Number   of Counties |
| --- | --- | --- | --- | --- | --- |
| *Mangifera indica* | Mango | Introduced | 38 | 44 | 13 |
| *Metopium toxiferum* | Poisonwood | Native | 96 | 117 | 6 |
| *Rhus copallina* | Winged Sumac | Native | 205 | 291 | 57 |
| *Schinus terebinthifolia* | Brazilian Pepper | Introduced,  Invasive | 322 | 493 | 35 |
| *Toxicodendron pubescens* | Eastern Poison Oak | Native | 30 | 36 | 17 |
| *Toxicodendron radicans* | Eastern Poison Ivy | Native | 268 | 378 | 58 |
| *Toxicodendron vernix* | Poison Sumac | Native | 34 | 46 | 21 |

Wunderlin, R. P., B. F. Hansen, A. R. Franck, and F. B. Essig. 2019. Atlas of Florida Plants. http://florida.plantatlas.usf.edu/.

Appendix B. Full Likelihoods

**Data Model**

$$y_{i,t} | Z_{i}\sim Bernoulli (p_{i,t}\cdot Z_{i})$$

**Process Model**

$$Z_{i}\sim Bernoulli\left( \psi_{i} \right)$$

$$logit \left( \psi_{i} \right)=\alpha_{0}+\alpha_{1}x_{1}+\alpha_{2}x_{2}+ \alpha_{3}x_{3}$$

$$logit\left( p \right)=\epsilon_{col}+\epsilon_{mos}+\epsilon_{yr}+ \beta_{1}x_{4}+\beta_{2}x_{5}$$

**Priors**

$$\alpha_{0}\sim Uniform\left( -5,5 \right)$$

$$\alpha_{1}\sim Uniform\left( -5,5 \right)$$

$$\alpha_{2}\sim Uniform\left( -5,5 \right)$$

$$\alpha_{3}\sim Uniform\left( -5,5 \right)$$

$$\beta_{1}\sim Uniform \left( -5,5 \right)$$

$$\beta_{2}\sim Uniform\left( -5,5 \right)$$

$$\epsilon_{col}\sim Normal\left( 0, \tau_{obs} \right)$$

$$\epsilon_{mos}\sim Normal\left( 0, \tau_{mos} \right)$$

$$\epsilon_{yr} \sim Normal\left( 0, \tau_{yr} \right)$$

**Hyperpriors**

$$\tau_{col}\sim Gamma \left( 1, 0.001 \right)$$

$$\tau_{mos}\sim Gamma\left( 1, 0.001 \right)$$

$$\tau_{yr} \sim Gamma(1, 0.001)$$

Appendix C: Species-specific collection biases

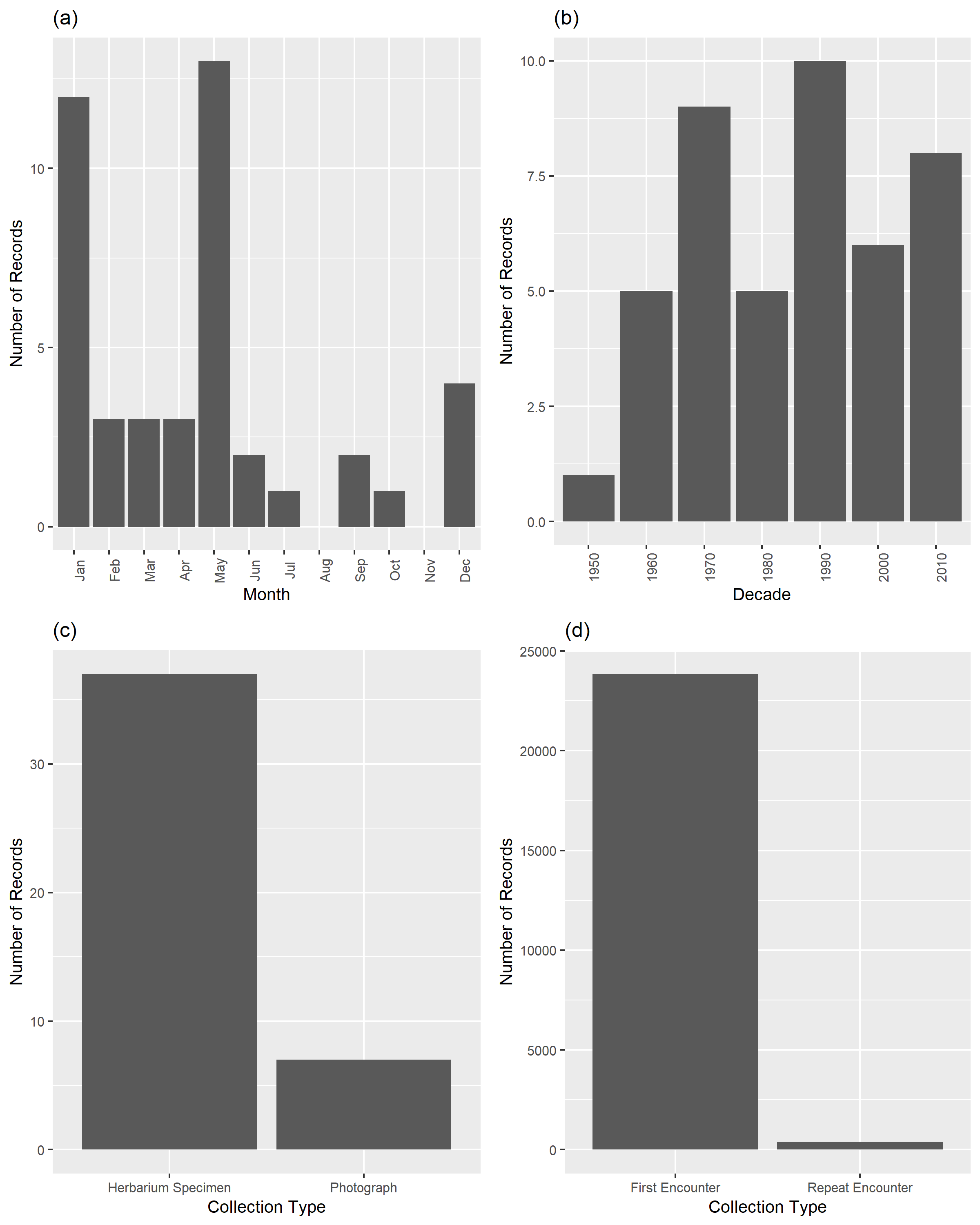

Figure C.1. (a) Number of records collected in each month of the year, (b) Number of records collected in each decade, (c) Number of records that are museum specimens or photographs and (d) Number of records that represent first encounters or repeat encounters of *M. indica.*

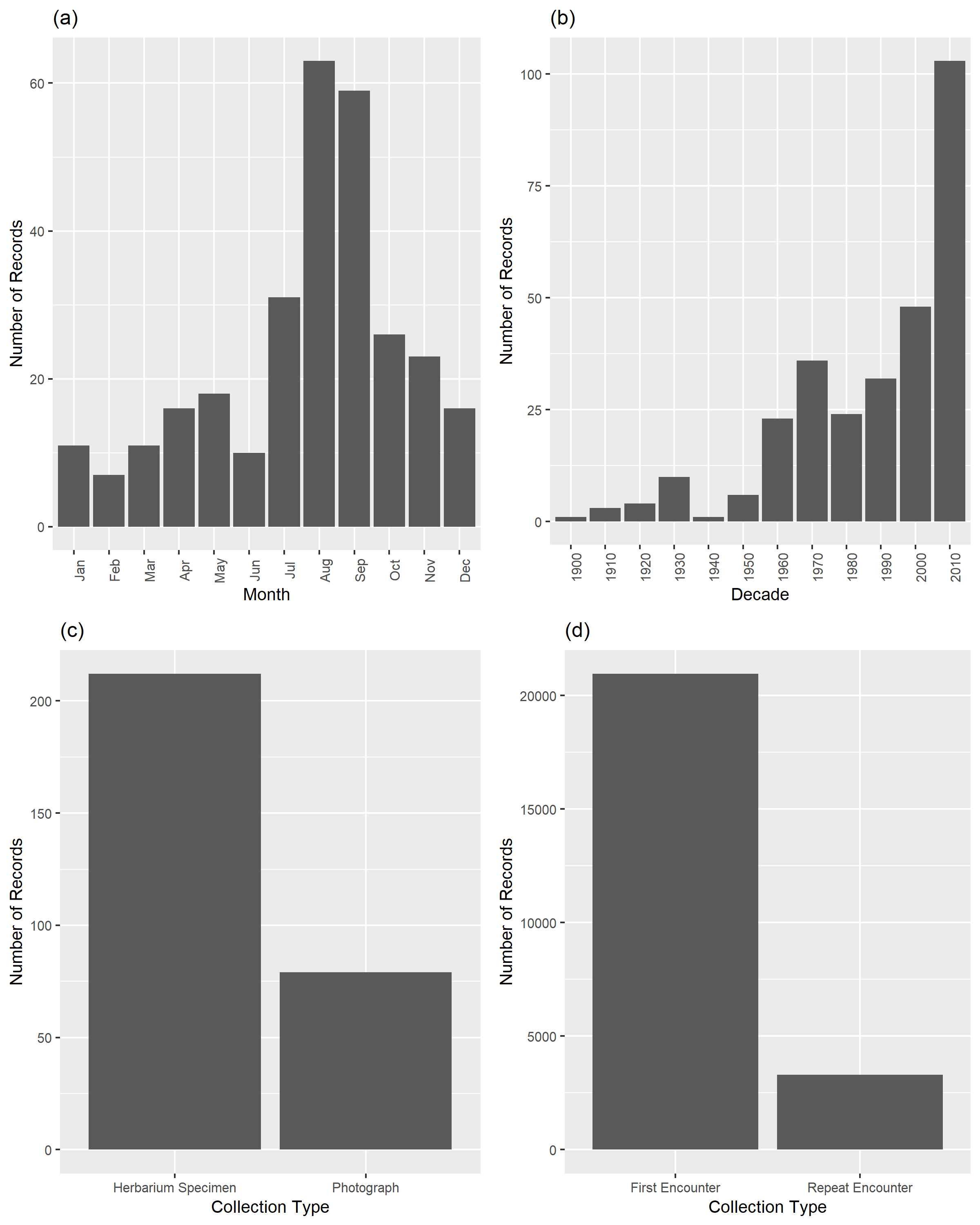

Figure C.2. (a) Number of records collected in each month of the year, (b) Number of records collected in each decade, (c) Number of records that are museum specimens or photographs and (d) Number of records that represent first encounters or repeat encounters of *R. copallina.*

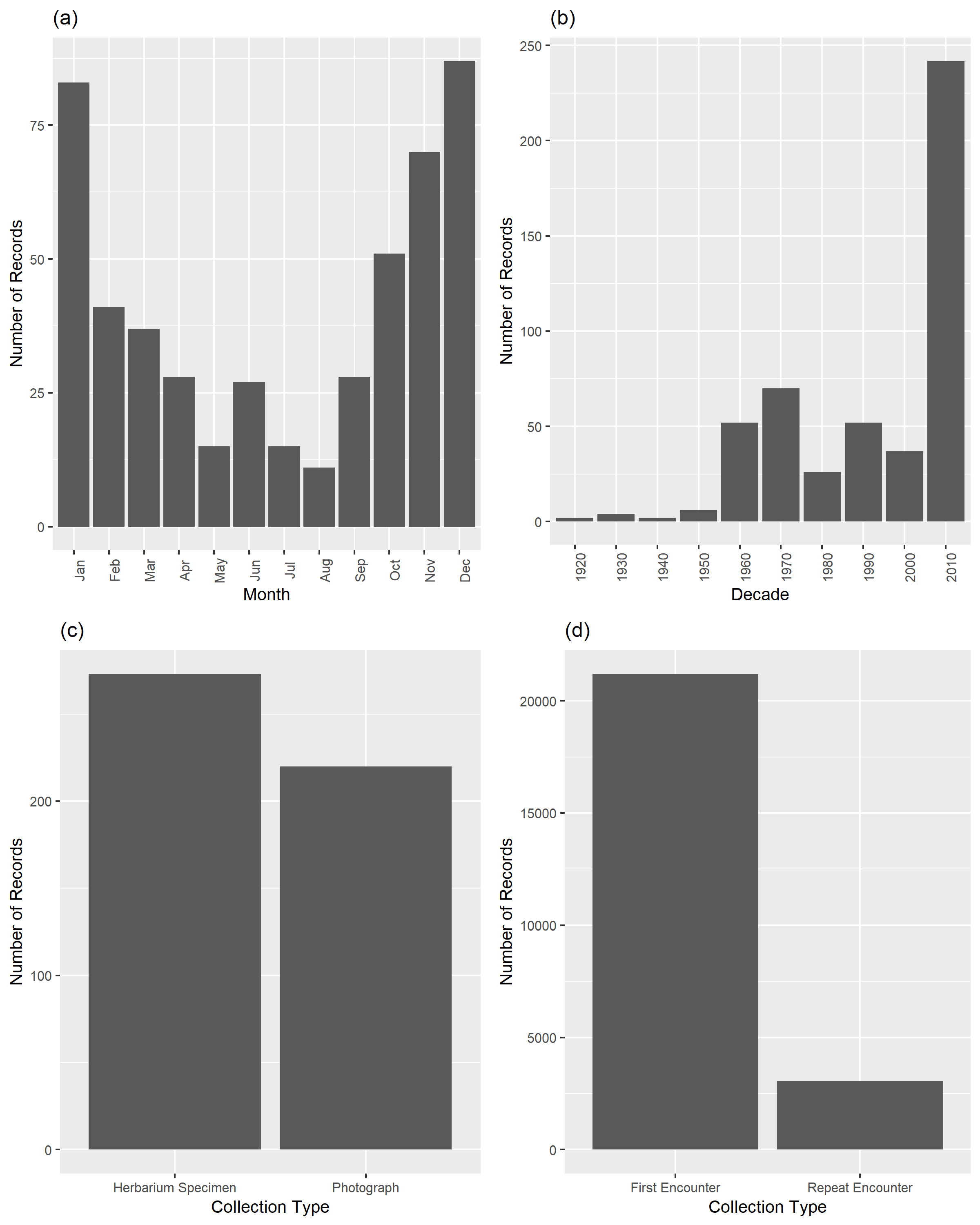

Figure C.3. (a) Number of records collected in each month of the year, (b) Number of records collected in each decade, (c) Number of records that are museum specimens or photographs and (d) Number of records that represent first encounters or repeat encounters of *S. terebinthifolia.*

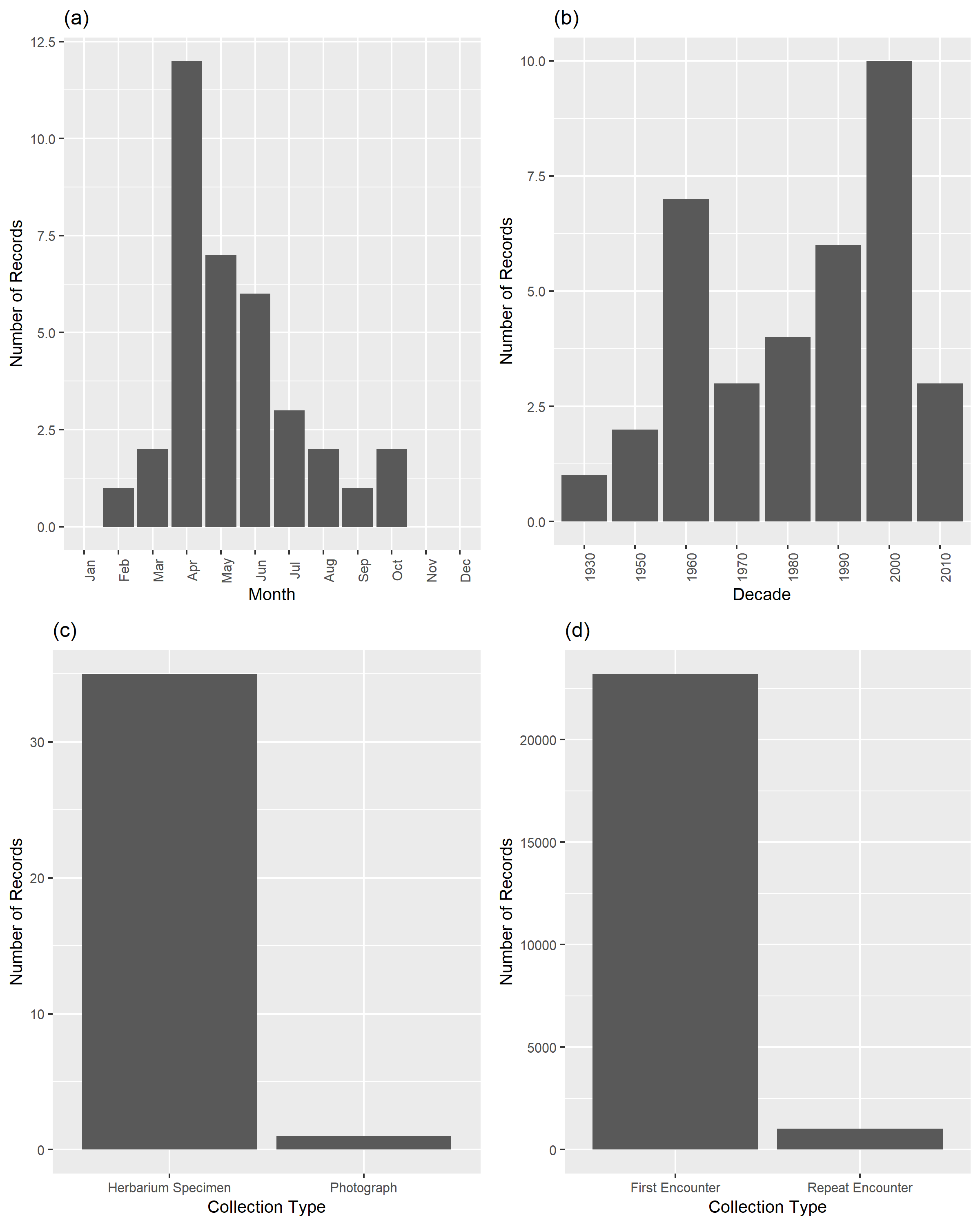

Figure C.4. (a) Number of records collected in each month of the year, (b) Number of records collected in each decade, (c) Number of records that are museum specimens or photographs and (d) Number of records that represent first encounters or repeat encounters of *T. pubescens.*

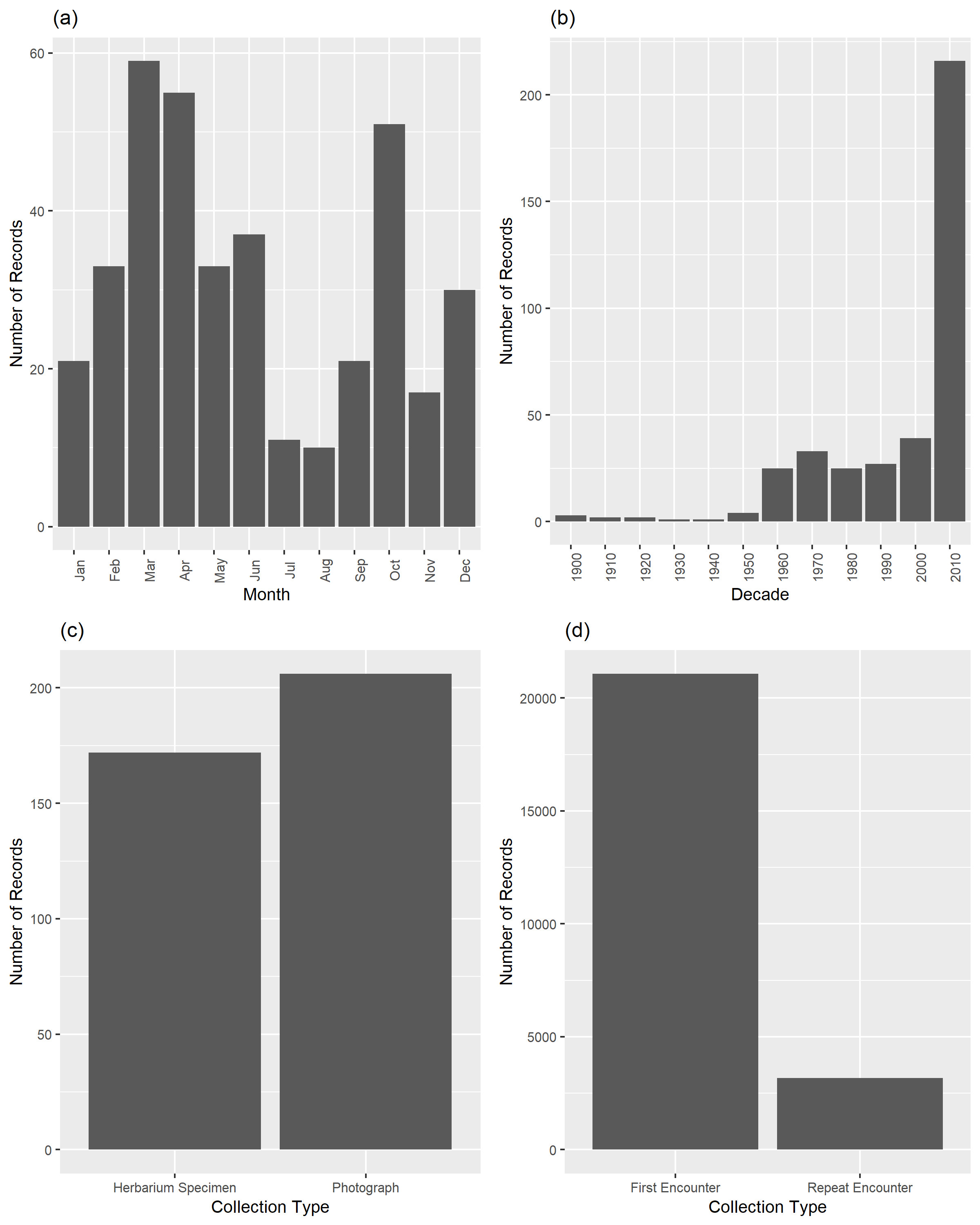

Figure C.5. (a) Number of records collected in each month of the year, (b) Number of records collected in each decade, (c) Number of records that are museum specimens or photographs and (d) Number of records that represent first encounters or repeat encounters of *T. radicans.*

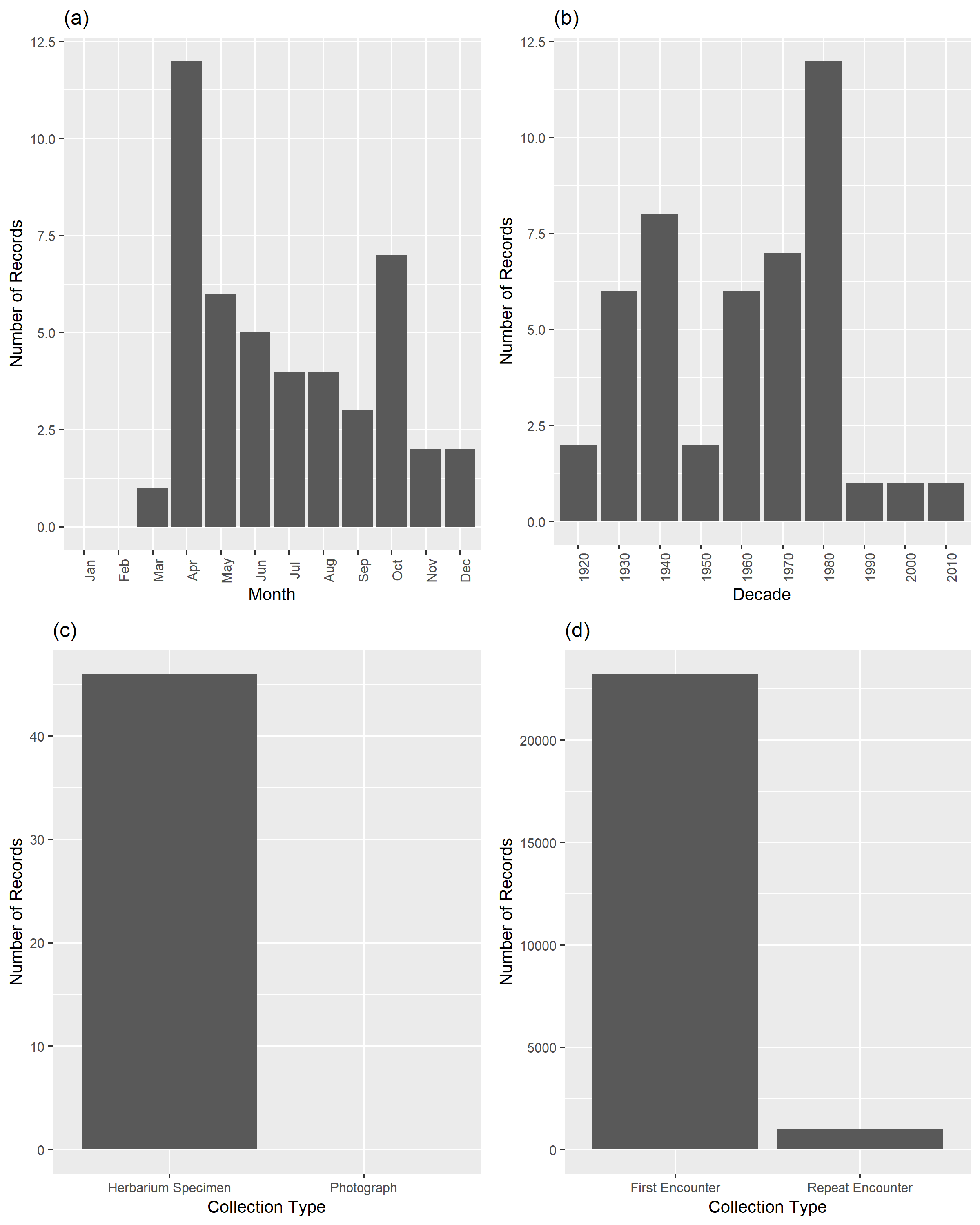

Figure C.6. (a) Number of records collected in each month of the year, (b) Number of records collected in each decade, (c) Number of records that are museum specimens or photographs and (d) Number of records that represent first encounters or repeat encounters of *T. vernix.*

**Appendix D: Posterior Caterplots of Occupancy Covariates**

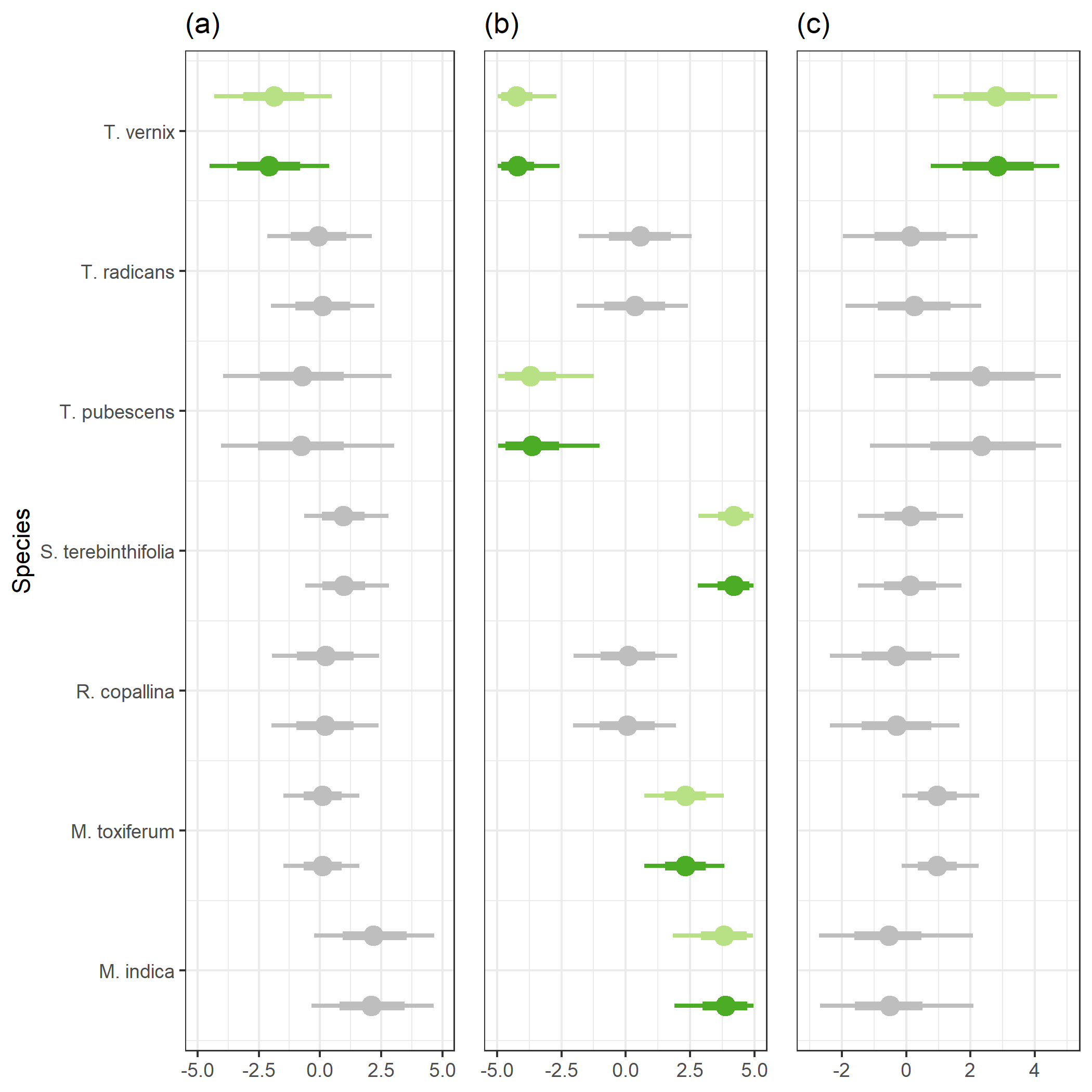

Figure D.1 50% (thin line) and 95% (thick line) posterior credible intervals for (a) human population density, (b) minimum temperature and (c) county area. For each species, the bottom line (dark green) is the credible interval for the parameter estimate from the full model with all data and the top line (light green) is the credible interval for the parameter estimate from the full model with filtered data. Grey lines indicate that the 95% credible interval overlaps with zero.

**Appendix E: Posterior caterplots of detection random effects for remaining species**

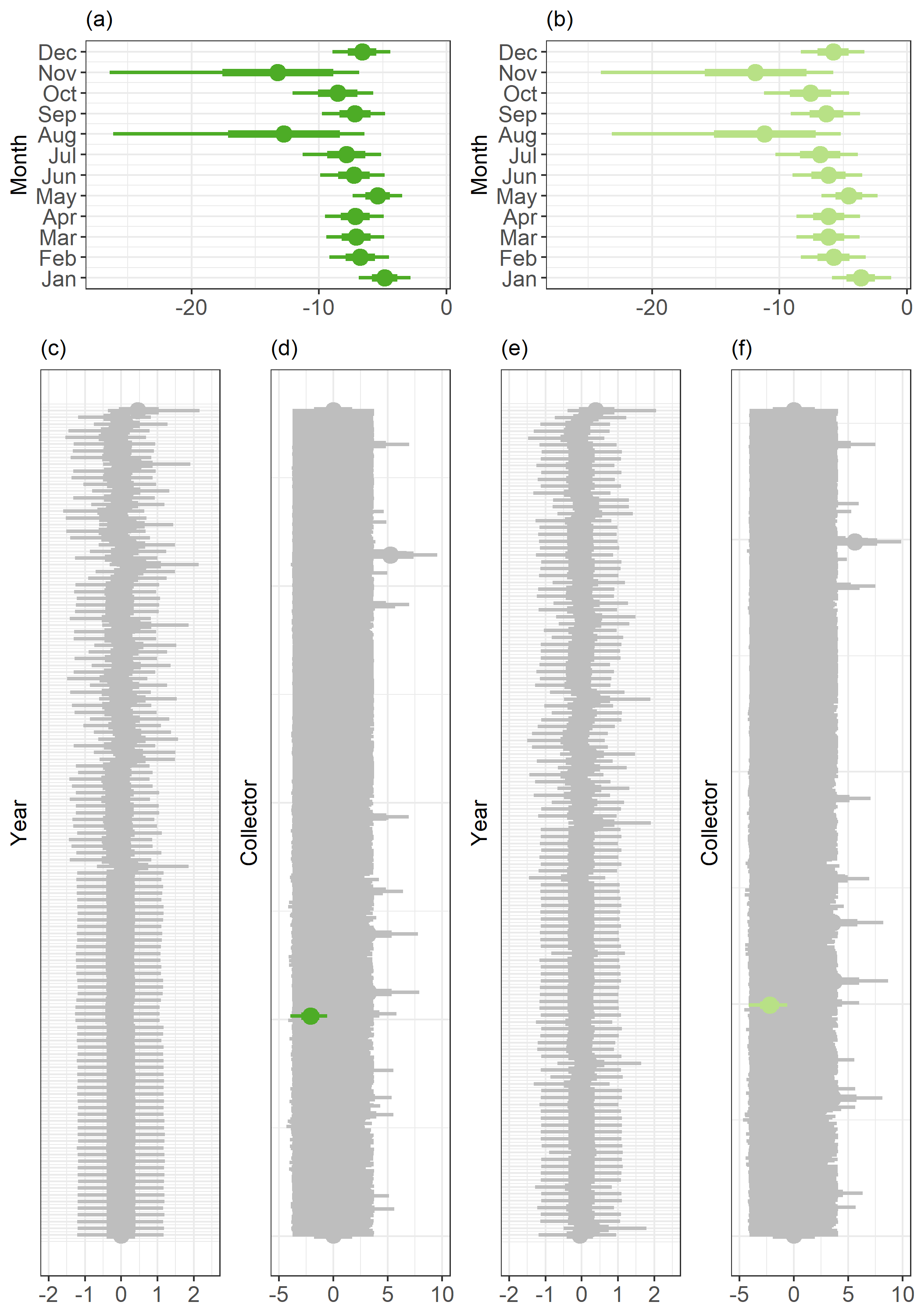

Figure E.1 *M. indica* 50% (thick line) and 95% (thin line) credible intervals for (a) month effect for the full model with all data, (b) month effect for the full model with filtered data, (c) collector effect for the full model with all data (d) year effect for the full model with all data (e) collector effect for the full model with filtered data, and (f) year effect for the full model with filtered data. Greyed-out credible intervals overlap 0.

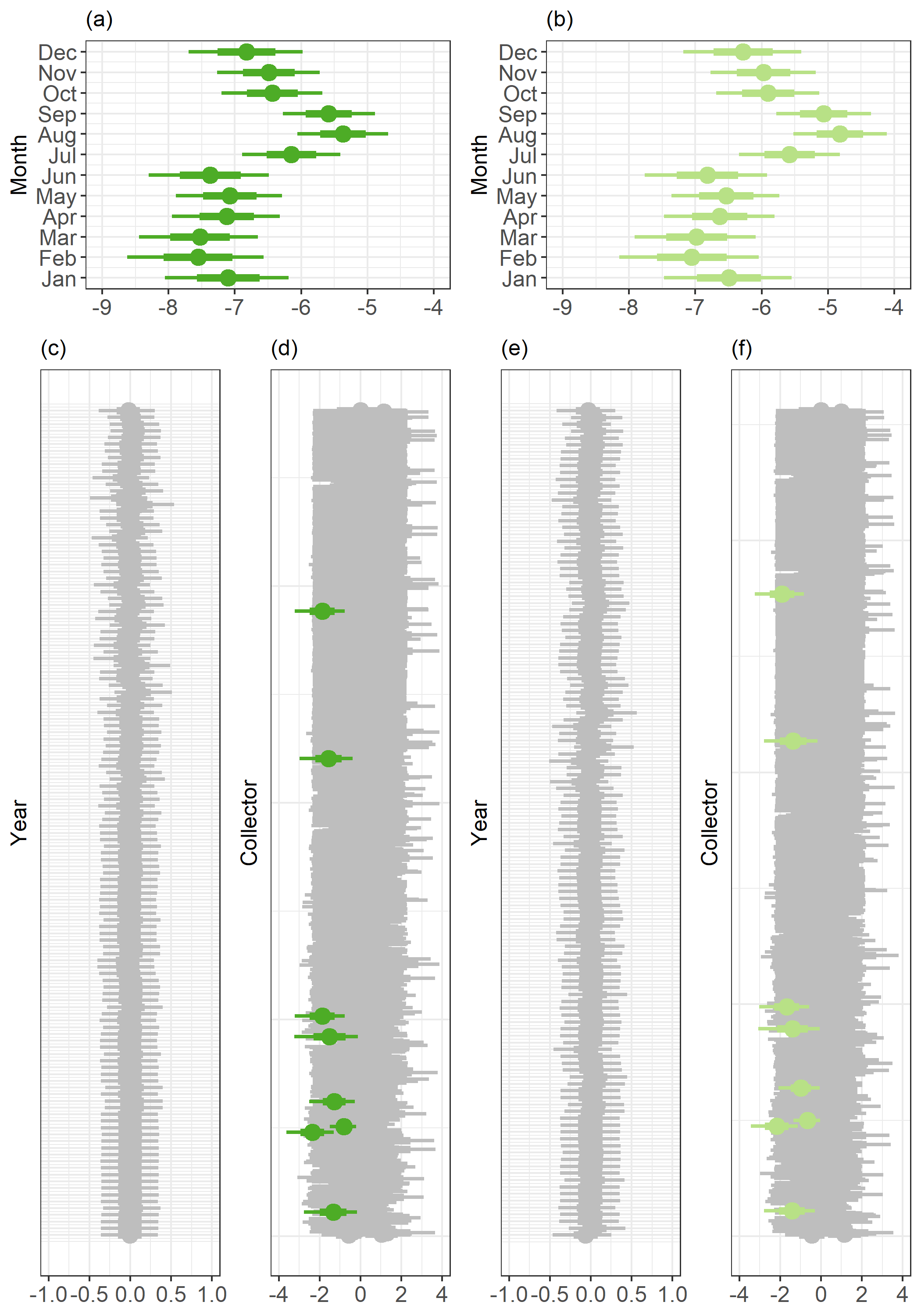

Figure E.2. *R. copallina* 50% (thick line) and 95% (thin line) credible intervals for (a) month effect for the full model with all data, (b) month effect for the full model with filtered data, (c) collector effect for the full model with all data (d) year effect for the full model with all data (e) collector effect for the full model with filtered data, and (f) year effect for the full model with filtered data. Greyed-out credible intervals overlap 0.

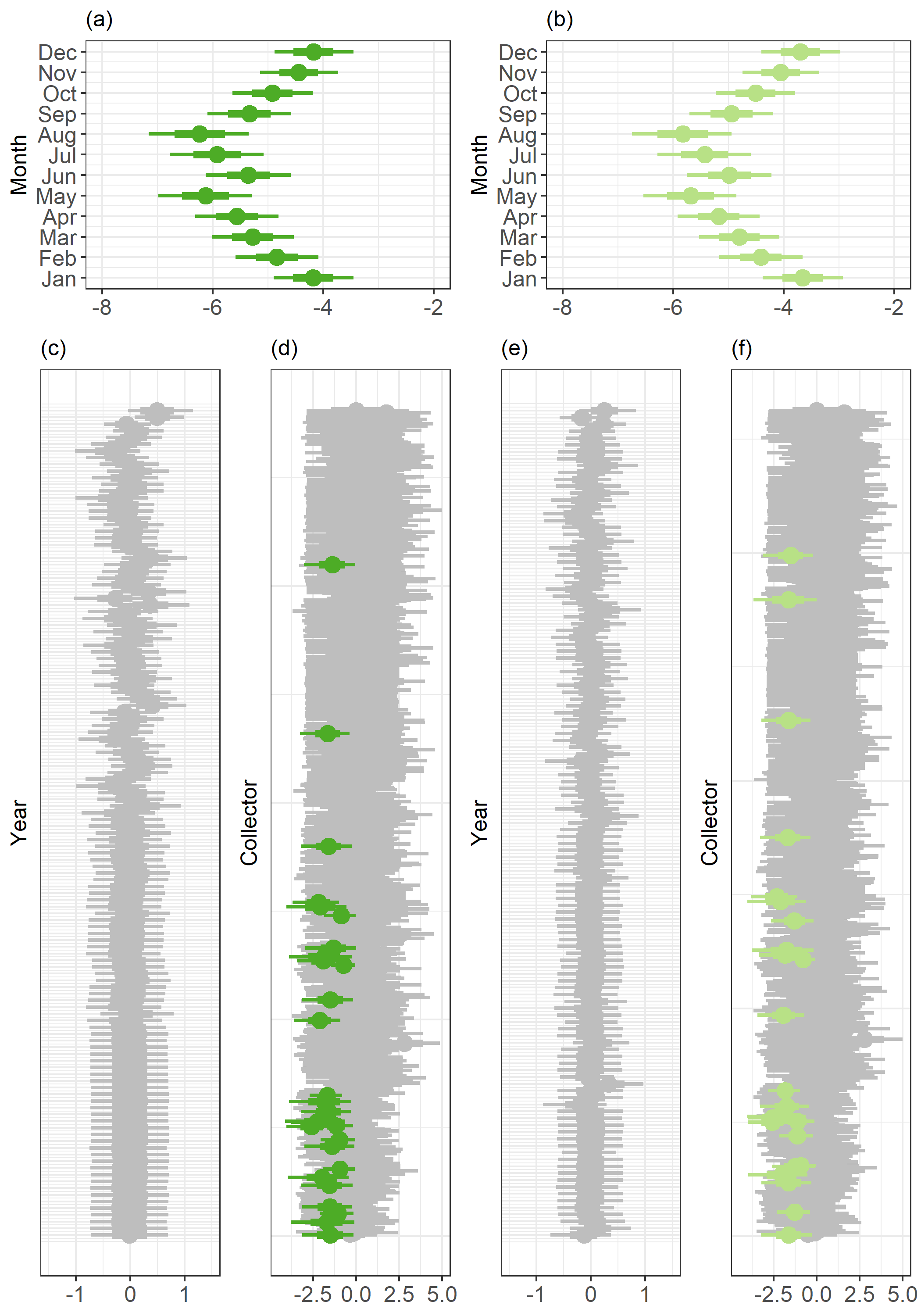
 Figure E.3. *S. terebinthifolia* 50% (thick line) and 95% (thin line) credible intervals for (a) month effect for the full model with all data, (b) month effect for the full model with filtered data, (c) collector effect for the full model with all data (d) year effect for the full model with all data (e) collector effect for the full model with filtered data, and (f) year effect for the full model with filtered data. Greyed-out credible intervals overlap 0.

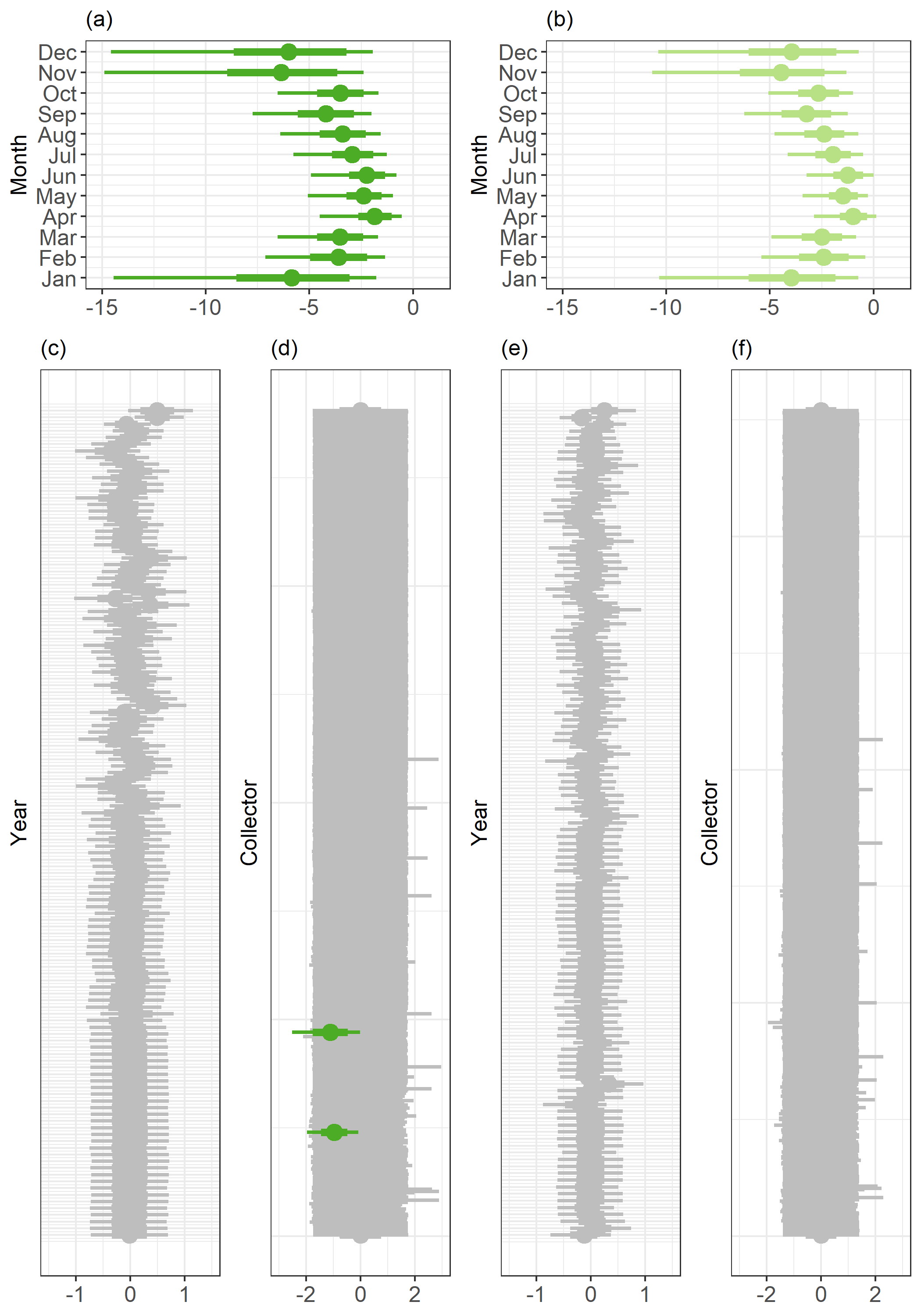

Figure E.4. *T. pubescens* 50% (thick line) and 95% (thin line) credible intervals for (a) month effect for the full model with all data, (b) month effect for the full model with filtered data, (c) collector effect for the full model with all data (d) year effect for the full model with all data (e) collector effect for the full model with filtered data, and (f) year effect for the full model with filtered data. Greyed-out credible intervals overlap 0.

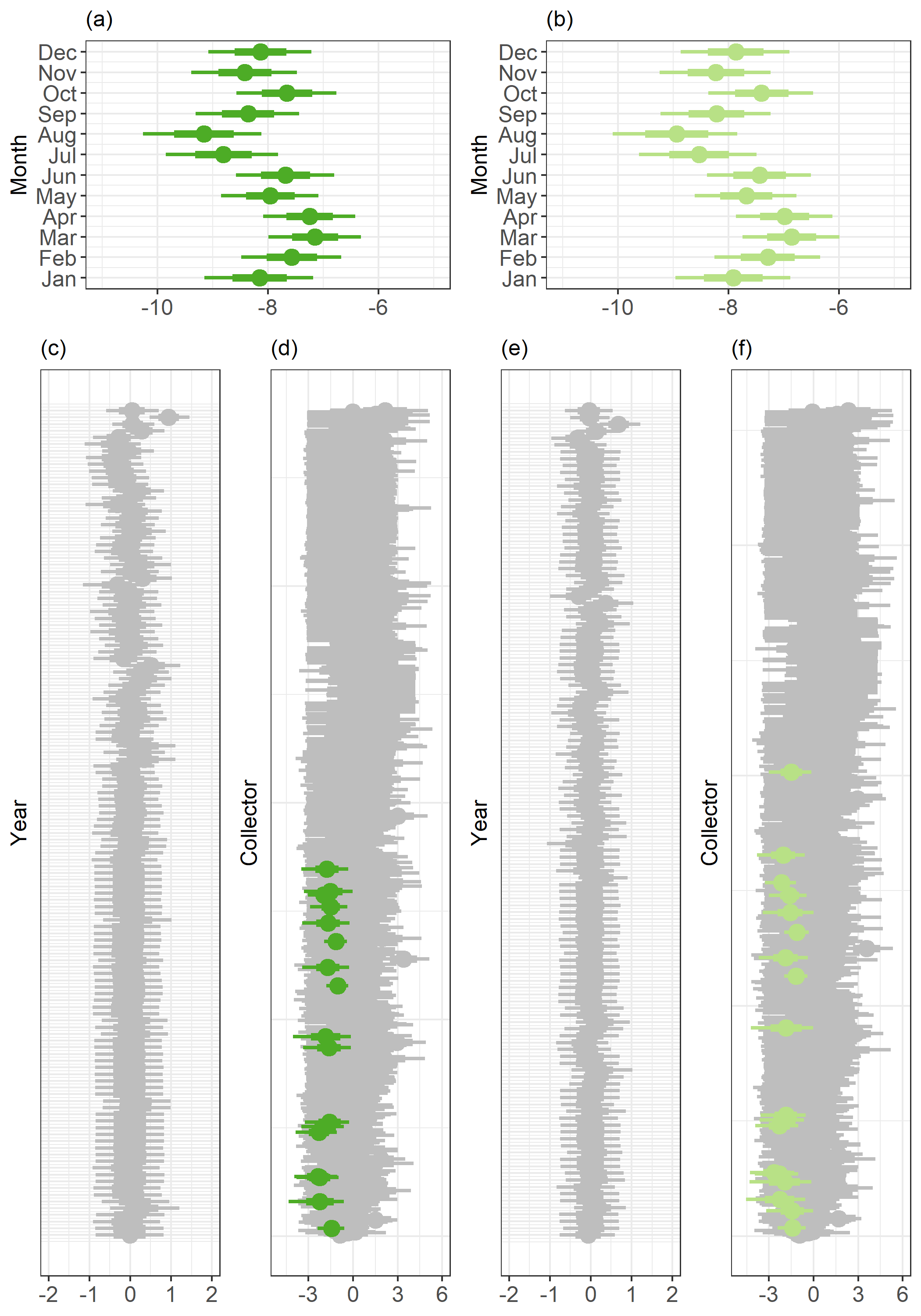

Figure E.5. *T. radicans* 50% (thick line) and 95% (thin line) credible intervals for (a) month effect for the full model with all data, (b) month effect for the full model with filtered data, (c) collector effect for the full model with all data (d) year effect for the full model with all data (e) collector effect for the full model with filtered data, and (f) year effect for the full model with filtered data. Greyed-out credible intervals overlap 0.

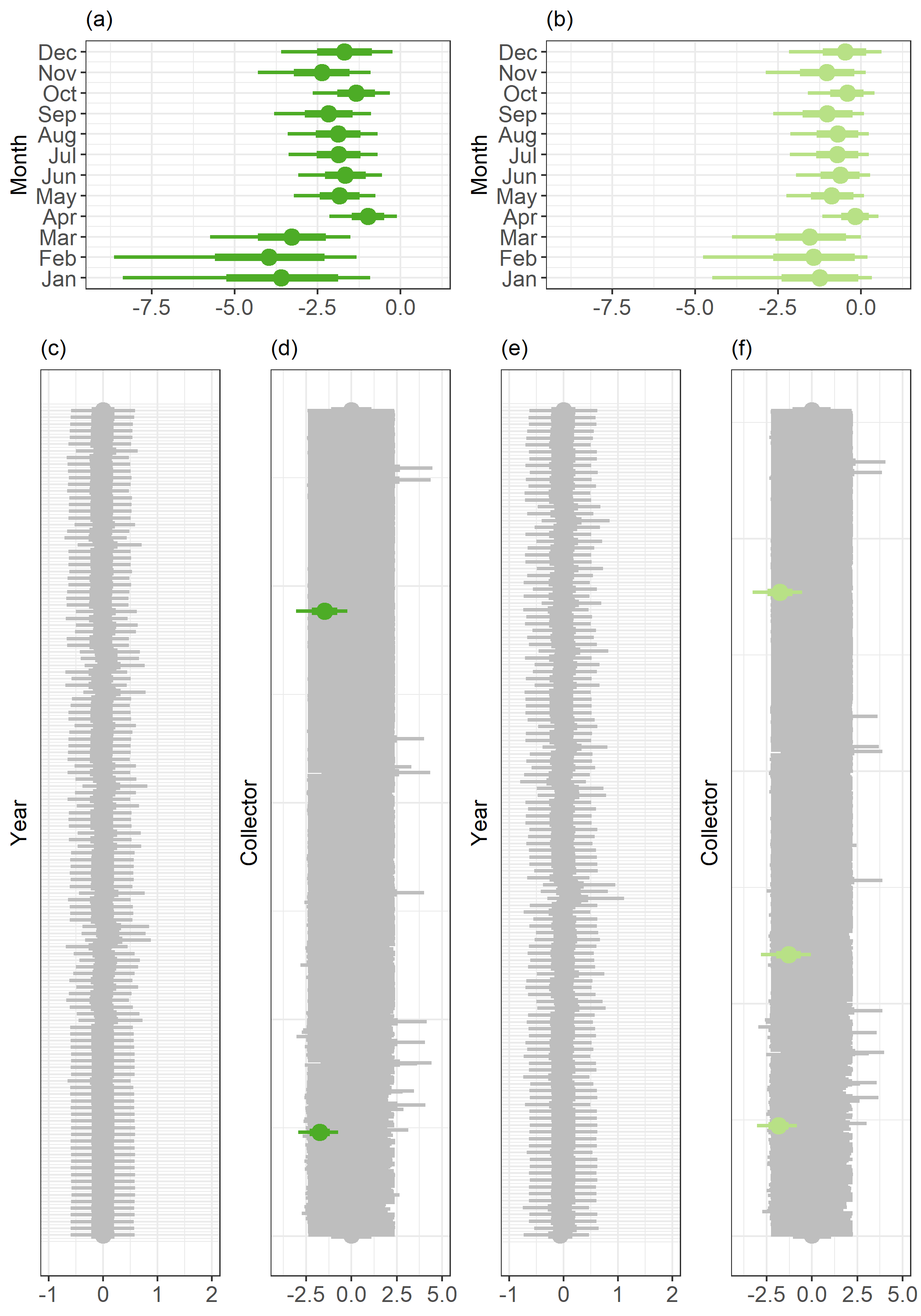

Figure E.6. *T. vernix* 50% (thick line) and 95% (thin line) credible intervals for (a) month effect for the full model with all data, (b) month effect for the full model with filtered data, (c) collector effect for the full model with all data (d) year effect for the full model with all data (e) collector effect for the full model with filtered data, and (f) year effect for the full model with filtered data. Greyed-out credible intervals overlap 0.
